# Bacterial FtsZ induces mitochondrial fission in human cells

**DOI:** 10.1101/2020.01.24.917146

**Authors:** Anna Spier, Martin Sachse, Nam To Tham, Mariette Matondo, Pascale Cossart, Fabrizia Stavru

**Affiliations:** Unité des Interactions Bactéries-Cellules, Institut Pasteur, Paris, France; Institut National de la Santé et de la Recherche Médicale (INSERM), U604, Paris, France; Institut National de la Recherche Agronomique (INRA), USC2020, Paris, France; Université Paris Diderot, Sorbonne Paris Cité, Paris, France; Unité Technologie et service BioImagerie Ultrastructurale, Institut Pasteur, Paris, France; Plateforme Protéomique, Unité de Spectrometrie de Masse pour Biologie (UTechS MSBio), Institut Pasteur, Paris, France; Centre National de la Recherche Scientifique (CNRS), USR 2000, Paris, France; CNRS SNC5101, Paris, France

**Keywords:** mitochondrial division, bacterial division, Drp1, mtDNA, inner mitochondrial membrane

## Abstract

Mitochondria are key eukaryotic organelles that evolved from an intracellular bacterium, in a process involving bacterial genome rearrangement and streamlining. As mitochondria cannot form *de novo*, their biogenesis relies on growth and division. In human cells, mitochondrial division plays an important role in processes as diverse as mtDNA distribution, mitochondrial transport and quality control. Consequently, defects in mitochondrial division have been associated with a wide range of human pathologies. While several protists have retained key components of the bacterial division machinery, none have been detected in human mitochondria, where the dynamin-related protein Drp1, a cytosolic GTPase is recruited to the mitochondrial outer membrane, forming helical oligomers that constrict and divide mitochondria. Here, we created a human codon optimized version of FtsZ, the central component of the bacterial division machinery, and fused it to a mitochondrial targeting sequence. Upon expression in human cells, mt-FtsZ was imported into the mitochondrial matrix, specifically localizing at fission sites prior to Drp1 and significantly increasing mitochondrial fission levels. Our data suggests that human mitochondria have an internal, matrix-localized fission machinery, whose structure is sufficiently conserved as to accommodate bacterial FtsZ. We identified interaction partners of mt-FtsZ, and show that expression of PGAM5, FAM210, SFXN3 and MTCH1 induced mitochondrial fission. Our results thus represent an innovative approach for the discovery of novel critical mitochondrial fission components.

## Introduction

Mitochondria are key eukaryotic organelles, which have retained their own genome and are delimited by two membranes. The bacterial origin of mitochondria originally proposed by Lynn Margulis (then Lynn Sagan^1^) is now largely accepted, and even complex mitochondrial features, such as the inner mitochondrial membrane invaginations termed cristae, have recently been shown to be evolutionarily conserved in specific bacterial lineages^2^. However, one conundrum is the apparent lack of evolutionary conservation of the division machinery between bacteria and mitochondria of fungi and metazoa.

Division is an essential process for both bacteria and mitochondria. In the vast majority of bacteria, cell division is performed by a multiprotein complex, at the heart of which lies the tubulin homologue and evolutionarily conserved protein FtsZ^3–5^. In bacteria, FtsZ assembles early at prospective fission sites, forming the Z-ring and recruiting several additional proteins to mediate cell division (reviewed in ^6,7^).

Mitochondrial fission is necessary for proper distribution of the organelle during mitosis and in highly polarized cells^8^ and the dynamic equilibrium between fission and fusion is tightly connected to mitochondrial function in both human and yeast cells^9^. At the molecular level, important differences exist between organisms. While several protists have retained key components of the bacterial division machinery, none have been detected in fungi and metazoa^10^, where mitochondrial fission is thought to be governed by a cytosolic machinery^11^. An intermediate situation has been described in the red alga *Cyanoschizon merolae*, where assembly of an intramitochondrial FtsZ-based fission machinery appears coordinated with the assembly of a cytosolic Dynamin-based fission machinery^12,13^. In human cells a series of events involving the ER, the actin and septin cytoskeleton and receptors on the mitochondrial outer membrane (OMM) culminates in the assembly of a dynamin-related protein (Drp1) on the OMM^8^; Drp1 then constricts mitochondria, leading to division, with potential synergistic action of Dyn2^14–16^. Fission ensues after constriction beyond a critical threshold^17^. During mitochondrial fission the two membranes that delimit mitochondria represent a challenge, as fusion between the outer and inner mitochondrial membranes has to be prevented to avoid leakage of mitochondrial content (Fig 1A). One possible scenario is that the inner membranes reach the necessary curvature and fusogenic distance earlier than the outer membrane, spontaneously fuse and retract, leaving only the outer membranes to fuse, leading to abscission of the two daughter mitochondria. Other scenarios invoke the presence of molecular machineries that either “insulate” the outer from the inner membrane during fission, or that specifically promote fission of the matrix compartment. The latter hypothesis appears the most likely, as a matrix-localized, “bacteria-like” fission machinery with homologs of the bacterial division protein FtsZ at its core has been identified in several protists^10,18–21^. In light of the monophyletic origin of mitochondria, we and others postulated that metazoan mitochondria would also harbour a fission machinery located in the matrix^8,22^. Indeed, fission of the matrix compartment has been observed in the absence of outer membrane fission in several metazoans^23–26^. However, previous attempts at bioinformatic identification of a bacteria-derived division machinery in metazoan mitochondria based on sequence similarity have failed^10,27^. We hypothesized that the internal fission machinery of mitochondria might nevertheless have retained a certain degree of structural conservation with respect to its bacterial ancestor. To test this hypothesis, we asked whether the key orchestrator of bacterial cell division FtsZ would be able to induce mitochondrial fission in mammalian cells. We engineered synthetic constructs that allowed targeting of bacterial FtsZ into the mitochondrial matrix and found that alphaproteobacterial FtsZ (mt-αFtsZ) specifically localized at mitochondrial fission sites and substantially increased mitochondrial fission levels. As several proteins concur to recruit FtsZ to the membrane in bacteria, we explored which mitochondrial proteins might play this role in our experimental system. Among the interaction partners of mt-αFtsZ that we identified, we tested five transmembrane mitochondrial proteins, four of which induce mitochondrial fission upon overexpression, potentially representing new players in mammalian mitochondrial dynamics.

**Fig 1:**
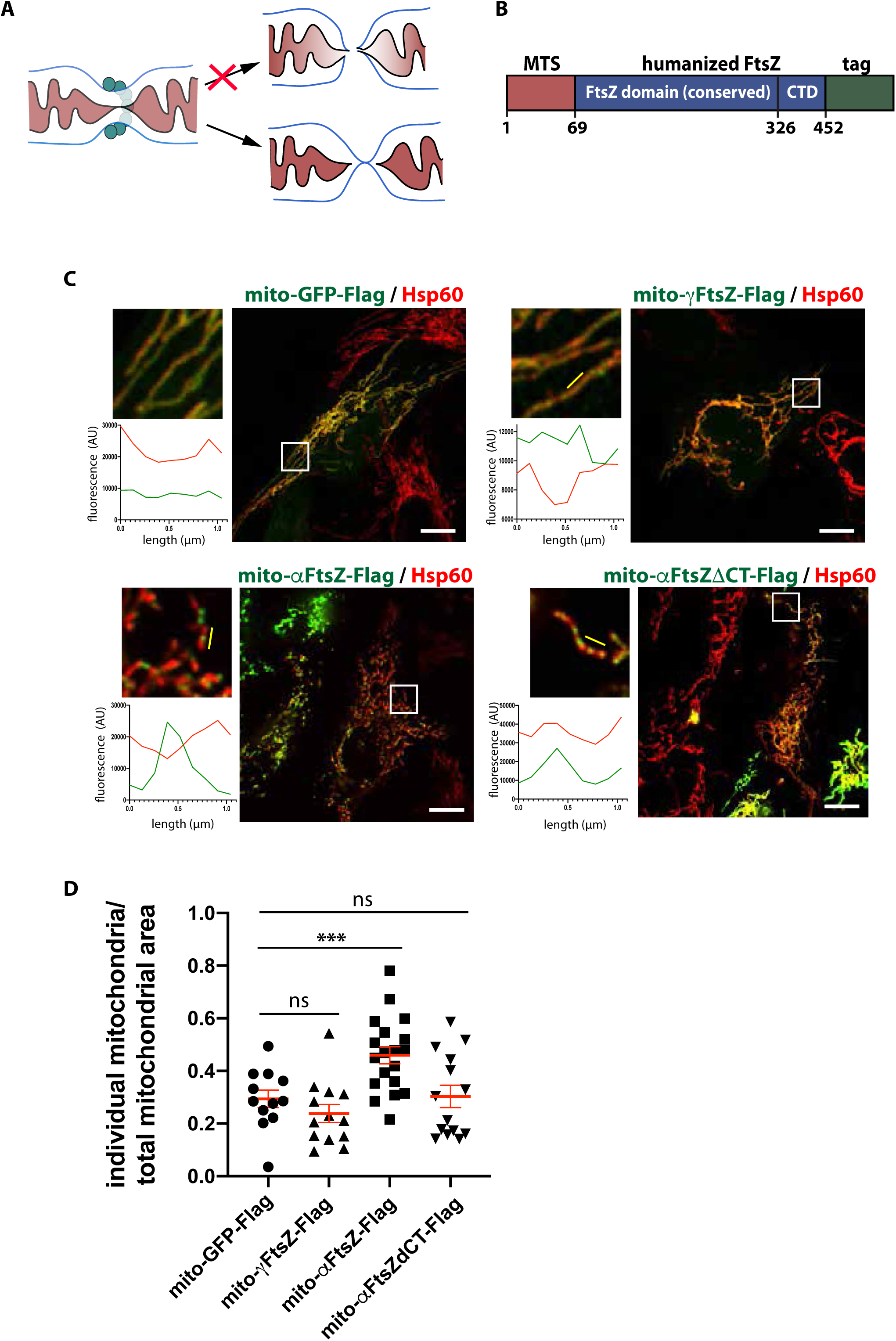
Expression of alphaproteobacterial FtsZ (mt-αFtsZ) in human mitochondria induces mitochondrial fission. **A** Schematic representation of mitochondrial constriction (cross-section), where Drp1 is depicted in turquoise, the outer membrane in blue, the inner membrane in black and the matrix in burgundy. The top arrow points at a theoretical scenario, where fission entails inner and outer membrane fusion. The bottom arrow points at the current fission model, where inner membrane undergoes homotypic fusion, leading to matrix fission. This is then followed by homotypic fusion of the outer membrane to achieve complete abscission of the two daughter mitochondria. *In vivo*, matrix and outer membrane fission appear closely linked in time and space. **B** General layout of the synthetic FtsZ constructs used in this study: a well-characterized mitochondrial targeting sequence (MTS) from subunit 9 of the Fo-ATPase of *N. crassa* was fused at the N-terminus of bacterial FtsZ, which was codon-optimized for expression in human cells. A C-terminal Flag or GFP tag was added for detection. Numbers refer to the alphaproteobacterial construct mt-αFtsZ. A control construct was created by replacing the FtsZ sequence with GFP, resulting in mito-GFP-Flag. **C** Immunofluorescence of U2OS cells transfected with mito-GFP-Flag or mitochondrially targeted FtsZ from a gamma- (mt-γFtsZ-Flag) or an alphaproteobacterium (mt-αFtsZ-Flag, C-terminal deletion mutant mt-αFtsZΔCT-Flag, revealed in green) and mitochondria (Hsp60, red). Scalebar: 10µm, insets are enlarged 4-fold. Linescan positions are indicated by a yellow line. **D** Semiautomated quantification (MiNA plugin) of mitochondrial morphology expressed by the amount of individual mitochondria/mitochondrial area in cells transfected as in B. Three experiments were pooled. Mean and SEM are displayed in red, P=0.0002 by one-way Anova, adjusted P-value = 0.0049.

## Results

### Mitochondrial expression of alphaproteobacterial FtsZ induces mitochondrial fission

To achieve fission of the mitochondrial matrix compartment, we hypothesized that mammalian mitochondria would contain a protein-based inner fission machinery (IFM). Given the endosymbiotic origin of mitochondria, we reasoned that the IFM would not have evolved *de novo* and even though not detectable by sequence similarity^10,27^, it would still share structural features with the bacterial fission machinery.

To test whether mitochondria contain an IFM in the matrix that can interact with a bacterial fission protein, we decided to transiently express bacterial FtsZ in human cell lines (U2OS and HeLa). To this end, we constructed synthetic versions of FtsZ, which were codon-optimized for expression in human cells and fused to an N-terminal mitochondrial targeting sequence to allow import into the mitochondrial matrix, and a C-terminal tag to allow detection (Fig 1B). Although mitochondria have long been thought to derive from alphaproteobacteria^28–33^, recent work suggests that mitochondria evolved from a proteobacterial lineage which branched off before the alphaproteobacteria^34^. As the precise bacterial lineage that gave rise to mitochondria remains a matter of debate, we chose to express synthetic versions of both gamma- and alphaproteobacterial FtsZ (*Escerichia coli* and typhus group *Rickettsia* respectively, referred to as mt-αFtsZ and mt-γFtsZ). Immunofluorescence analysis of flag-tagged mt-αFtsZ showed that it localized to mitochondria (Fig 1C) in virtually all transfected cells (99.5±1.7%, n=1469, N=6 independent experiments). In 0.5±1.6% mt-αFtsZ was expressed at very high levels, formed filaments that colocalized with microtubules (Suppl. Fig 1A) and displayed dramatic mitochondrial fragmentation and perinuclear aggregation (suppl Fig 1B). In cells where mt-αFtsZ localized to mitochondria, those with intermediate and low levels of expression allowed the detection of mt-αFtsZ punctae, which appeared to accumulate at mitochondrial matrix constrictions. In contrast, mt-γFtsZ or the control construct mt-GFP did not accumulate at constrictions and often displayed a more even staining (Fig 1C). We then tested a C-terminal deletion mutant of mt-αFtsZ (mt-αFtsZΔCT) and found that it was able to polymerize in some cells (Fig 1C, suppl Fig 1C), but failed to localize at constrictions, in agreement with previous findings showing that while C-terminal deletion mutants of FtsZ are able to polymerize in *E.coli*, they do not support bacterial division^35^.

Next, we asked whether the ultrastructure of mitochondrial constrictions was affected by mt-αFtsZ. To do so we combined light microscopy with high pressure freezing electron microscopy in a correlative approach allowing us to focus on cells with intermediate expression levels. Mitochondrial constrictions did not qualitatively differ between mt-GFP and mt-αFtsZ expressing mitochondria, which displayed an inner diameter of 41.6nm versus 40.8nm respectively and an outer diameter of 57.1nm versus 62.7nm (suppl Fig 1D).

Given that mt-αFtsZ was found at constriction sites, we analysed whether it would induce mitochondrial fission by quantifying mitochondrial morphology with the semi-automatic ImageJ plugin MiNA. This analysis showed that full length mt-αFtsZ induced mitochondrial fission (Fig 1C, suppl Fig 1E). We validated our results by using a different cell line (HeLa) and manually measuring mitochondrial length (suppl. Fig 1F), which revealed a dose-dependent effect on mitochondrial fission. In addition, the slightly thicker mitochondria of HeLa cells allowed us to discern mt-αFtsZ-labelled matrix constrictions with non-constricted outer membrane, supporting previous reports showing that matrix constriction can occur in the absence of outer membrane constriction^25,26^.

These data show that in contrast to mt-γFtsZ, mt-αFtsZ specifically localizes to mitochondrial matrix constrictions and affects mitochondrial morphology, suggesting that it labels mitochondrial matrix fission sites.

### mt-αFtsZ localizes at mitochondrial fission sites prior to Drp1 recruitment

To assess whether mt-αFtsZ labeled constrictions indeed proceed to complete abscission, we followed GFP tagged mt-αFtsZ in mitochondria of live cells. Full length mt-αFtsZ labeled the vast majority of all fissions we observed (86.3%, n=73, N=6), accumulating at prospective fission sites and often distributing to the tips of both daughter mitochondria upon abscission (Fig 2A). In contrast, the C-terminal deletion mutant mt-αFtsZΔCT did not consistently label fission sites (Fig 2B) and was found at up to 1.3µm from the fission site in 8 out of 11 fissions (N=4). In agreement with these findings, mt-αFtsZΔCT displayed an almost two-fold decrease in inner mitochondrial membrane localization compared to full-length mt-αFtsZ as assessed by immune-electron microscopy (Fig 2C/D). In addition, we noticed that mt-αFtsZ induced a zipper-like phenotype with regular, closely apposed cristae, which we also observed by high pressure freezing electron microscopy.

**Fig 2:**
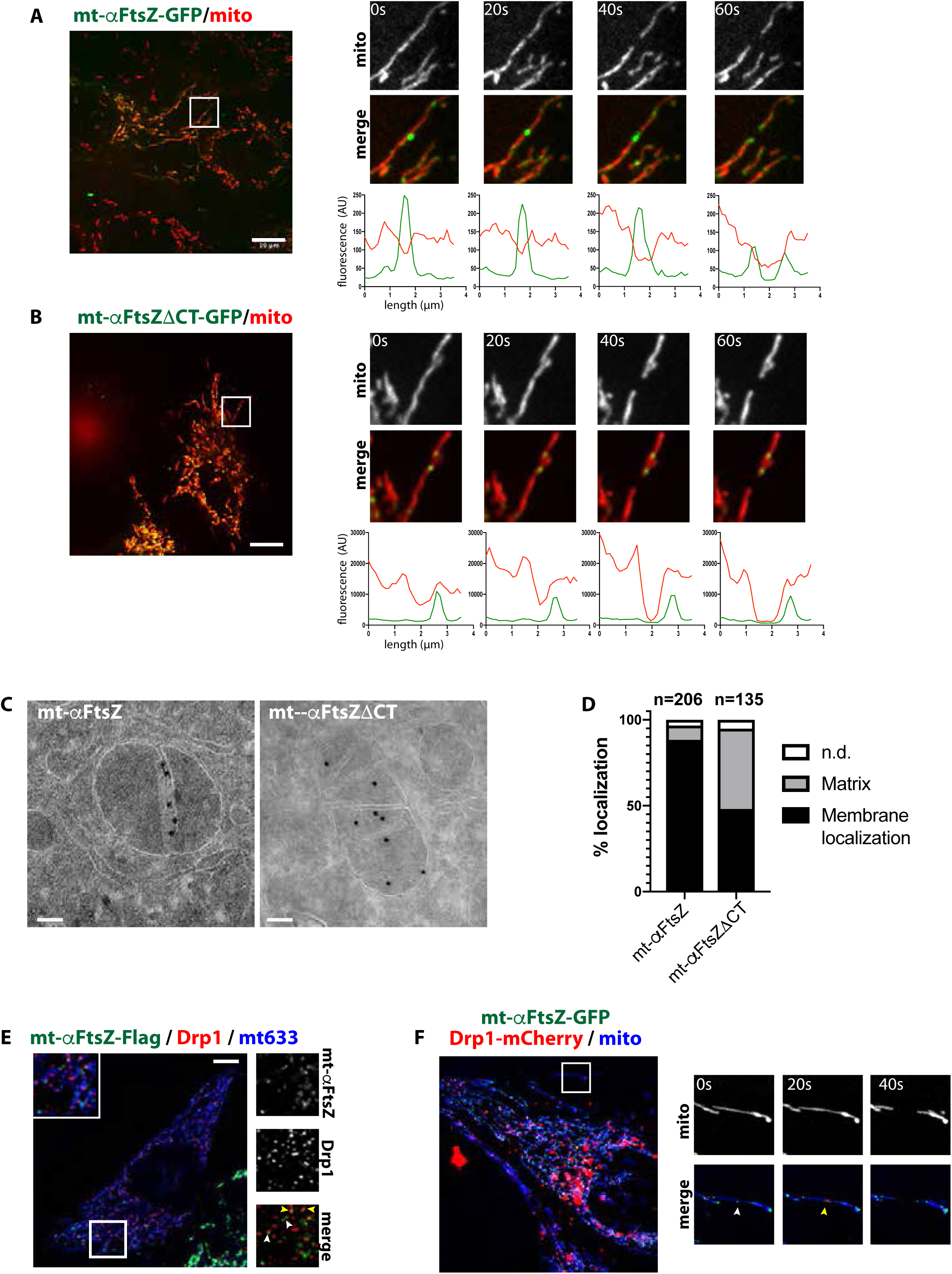
mt-αFtsZ labels mitochondrial fission sites and precedes Drp1 recruitment. **A** Live cell imaging of U2OS cells transfected with mt-αFtsZ-GFP or mt-αFtsZΔCT-GFP. Insets show an example of mitochondrial fission and are enlarged 2x. Linescans were taken for each timepoint along the fission axis. Scalebar: 10µm. **B** Post-embedding immuno-EM of HeLa cells transfected with the above constructs, stained with nanogold-anti-GFP. Scalebar: 100 nm. **C** Quantification of nanogold signal with respect to the inner mitochondrial membrane or the matrix (i.e. >15 nm distance from the inner membrane). n indicates the number of analyzed nanogold grains from 36 (mt-αFtsZ-GFP) or 25 random sections (mt-αFtsZΔCT-GFP). **D** Colocalization of mt-αFtsZ-GFP (green) with Drp1 (red) in Hela cells. Mitochondria were stained with mitotracker deep red (blue). Yellow arrowheads point at colocalization between mt-αFtsZ-GFP and Drp1, white arrowheads point at apposition. Inset enlarged 2x. **E** Live cell imaging of U2OS cells co-transfected with mt-αFtsZ-GFP and Drp1-mCherry, stained with mitotracker deep red. Insets show an example of mitochondrial fission and are enlarged 2x. mt-αFtsZ-GFP (white arrowhead) is present prior to Drp1 (yellow arrowhead).

Next, we investigated the spatiotemporal relationship between mt-αFtsZ and Drp1. A fraction of endogenous Drp1 colocalized with flag-tagged mt-αFtsZ or accumulated in its close proximity in fixed cells (Fig 2E, yellow and white arrowheads respectively). Live cell imaging revealed that mt-αFtsZ precedes Drp1 recruitment during mitochondrial fission (Fig 2F), suggesting that matrix constriction occurs prior to outer membrane constriction by Drp1. In agreement with this hypothesis, we found matrix constrictions that were labelled with mt-αFtsZ in the absence of outer membrane constriction (suppl Fig 2).

Toghether, these data indicate that matrix-localized mt-αFtsZ is a *bona fide* marker for mitochondrial fission sites and supports a fission model in which matrix constriction precedes Drp1 recruitment and mitochondrial abscission.

### Replication of the mitochondrial nucleoid is not necessary for mt-αFtsZ localization

The punctate pattern of mt-αFtsZ localization is reminiscent of nucleoids, which have been shown to accumulate at the tips of mitochondria^36,37^. We therefore assessed the spatial organization of mt-αFtsZ relative to nucleoids and found that mt-αFtsZ was excluded from the area occupied by nucleoids (Fig 3A), in particular at mitochondrial constrictions (suppl Fig 3). However, we also detected instances where mt-αFtsZ partially colocalized with nucleoids (Fig 3A, arrowheads). We hypothesized that this subset could represent replicating nucleoids, which have been estimated to amount to 9% of the total nucleoid population^38^ and to mark fission sites in yeast and human^39,40^. We thus examined whether mt-αFtsZ would colocalize with a red version of the mitochondrial DNA polymerase processivity subunit 2 (POLG2-mScarlet), which labels actively replicating mtDNA^39^. Surprisingly, mt-αFtsZ and POLG2 did not appear to substantially colocalize (Fig 3B).

**Fig 3:**
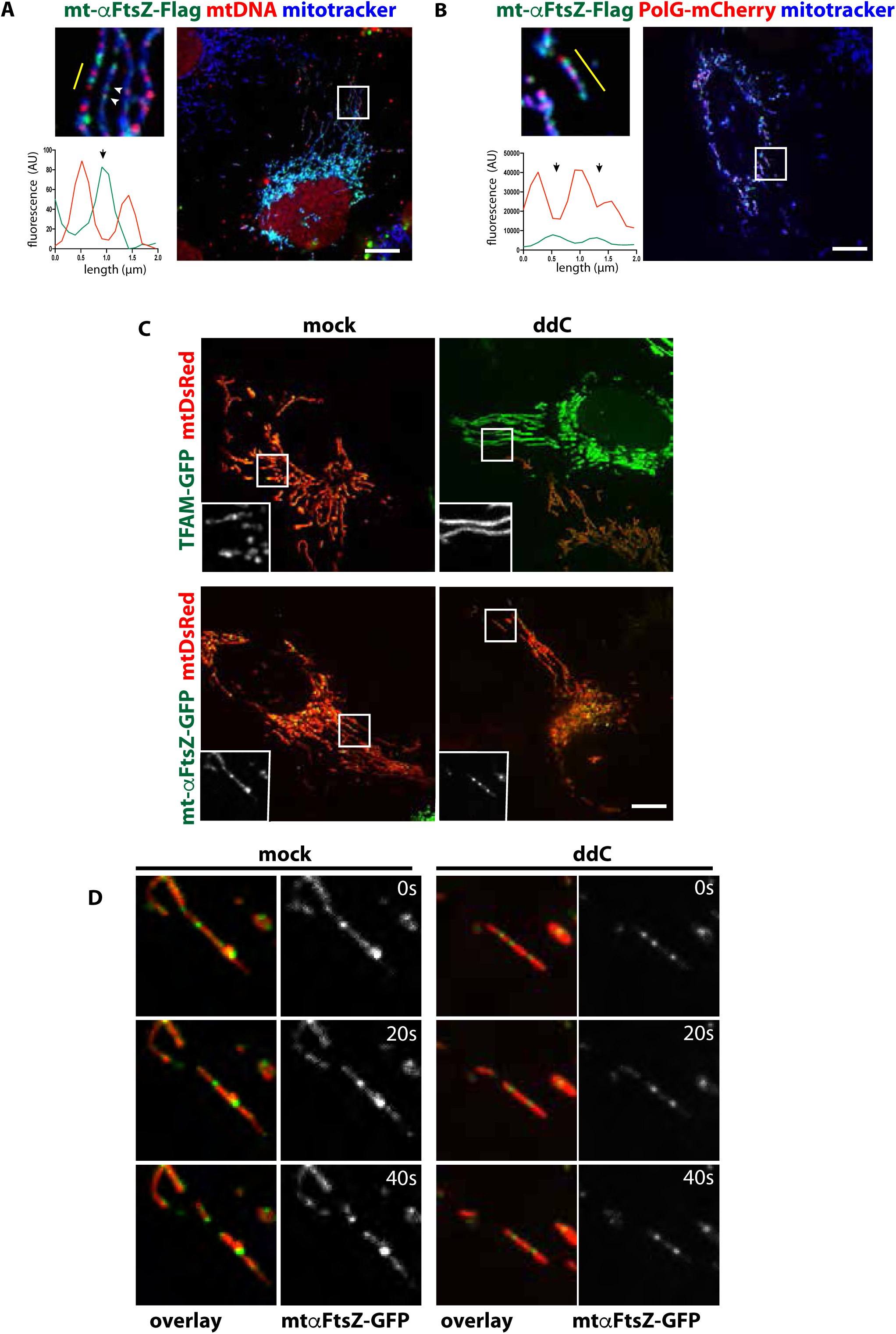
mt-αFtsZ localizes in proximity of the nucleoid, but is independent of mtDNA replication. **A** U2OS cells transfected with mt-αFtsZ-GFP (green) and labeled for mtDNA (red) and mitotracker deep red (blue). Inset enlarged 2x. Linescan showing juxtaposition of green and red signal; arrowheads point at colocalization beteeen mt-αFtsZ and mtDNA. Scalebar: 10µm. **B** Fluorescence images of U2OS cells transfected with mt-αFtsZ-GFP (green) and POLG-mScarlet (red). Mitochondria are shown in blue (mitotracker deep red). Inset enlarged 2x. Linescan showing alternating green and red signal. Scalebar: 10µm. **C** U2OS cells treated with 10µM ddC or vehicle for 48h, then transfected with mtDsRed and mt-αFtsZ-GFP or TFAM-GFP and imaged 36h later. Still images show diffuse staining of TFAM-GFP upon ddC treatment, indicating relocalization. mt-αFtsZ-GFP forms puncta irrespective of ddC treatment. Inset enlarged 2x. Scalebar: 10µm. **D** Time lapse imaging of the same cells depicted in C, showing 4x enlarged insets with mt-αFtsZ-GFP localization at fission sites in both control and ddC treated cells.

This prompted us to ask whether nucleoid replication was necessary for mt-αFtsZ localization and/or fission. We inhibited mtDNA replication with dideoxycytosine (ddC^41^). ddC treatment caused the nucleoid packing protein TFAM-GFP to label the entire matrix compartment, reflecting mtDNA depletion (Fig 3C). In contrast, mt-αFtsZ localization was not affected, marking mitochondrial fission sites in ddC treated and control cells (Fig3C/D). Together, these data strongly suggest that mt-αFtsZ localization is not dependent on nucleoid replication.

### Identification of mitochondrial proteins that interact with mt-αFtsZ

The dynamic localization of mt-αFtsZ during mitochondrial fission suggests that it interacts with a matrix-localized mitochondrial fission machinery. We therefore sought to identify interaction partners of mt-αFtsZ. To this end, we immunoprecipitated Flag-tagged mt-αFtsZ or mt-GFP from transiently transfected HeLa cells in presence of non-ionic detergent (0.5% NP40), and identified co-precipitating proteins by quantitative mass spectrometry. We identified 941 proteins, which only coprecipitated with mt-αFtsZ, but not with mt-GFP. In addition, 119 proteins were detected in both samples, but were significantly enriched in mt-αFtsZ immunoprecipitates (Fig 4A). 75% of the proteins we identified were not mitochondrial, reflecting mt-αFtsZ interactions taking place prior to its mitochondrial import and cells where mt-αFtsZ mislocalized to the cytoplasm due to high overexpression. As mislocalized mt-αFtsZ colocalizes with microtubules (suppl Fig 1A/B), we were not surprised to detect numerous proteins associated with microtubules among our hits. Interestingly, previous experiments have shown that *E.coli* FtsZ expressed in the cytosol of mammalian cells does not spontaneously colocalize with tubulin^42^, suggesting important structural differences between alpha- and gammaproteobacterial FtsZ. Among the overall 1060 interactants of mt-αFtsZ, 31.7% were organellar proteins and 25% (i.e. 269) mitochondrial according to the mitochondrial protein database IMPI (Integrated Mitochondrial Protein Index, v2018_Q2), despite the fact that we did not purify mitochondria prior to immunoprecipitation, reflecting an ∼3 fold enrichment in mitochondrial proteins if compared with ∼8% mitochondrial proteins in the human genome^43,44^. Consistently, Gene Ontology term analysis showed significant enrichment of organellar proteins, and in particular mitochondrial proteins (Fig 4B and suppl fig 4). Our dataset contained several inner and outer mitochondrial membrane proteins that have been linked to mitochondrial morphology or division (e.g. MICOS complex proteins Mic60, Mic27 and Mic19^45,46^, Prohibitin 2 and SPY complex members Yme1L and Stomatin-like protein 2^47^, mitochondrial fission protein 1 (Fis1^48^), mitochondrial fission process protein 1 (MTFP1/MTP18^49^), ATAD3B^50^, SLC25A46A^1^ and AFG3L2^51^, reinforcing our hypothesis that mt-aFtsZ can interact with an endogenous mitochondrial matrix fission machinery. Gene ontology analysis with the DAVID software highlighted an enrichment in mitochondrial inner membrane proteins (suppl fig 4A). Indeed, 74 of the 269 mitochondrial proteins (27.5%) we identified were predicted to contain at least one transmembrane domain. Interestingly, previous *in vitro* reconstitution experiments have shown that FtsZ cannot mediate unilamellar liposome constriction without its membrane-anchoring partner FtsA^52^. We therefore chose to focus on five highly enriched transmembrane proteins, the mitochondrial serine-threonine phosphatase PGAM5, MTCH1 (mitochondrial carrier homolog 1), FAM210A, the ATP synthase membrane subunit DAPIT (diabetes-associated protein in insulin-sensitive tissue) and SFXN3 (Sideroflexin 3). We selected these proteins based on their enrichment score and their detection in independent immunoprecipitation experiments, where beads were used as a control (not shown). Among the selected candidates, MTCH1, DAPIT and FAM210A have not been studied in the context of mitochondrial dynamics, while recent data has implicated PGAM5 in mitochondrial dynamics^53–55^ and in the course of this study members of the Sideroflexin family have been shown to act as serine transporters and impact mitochondrial morphology^56^.

**Fig 4:**
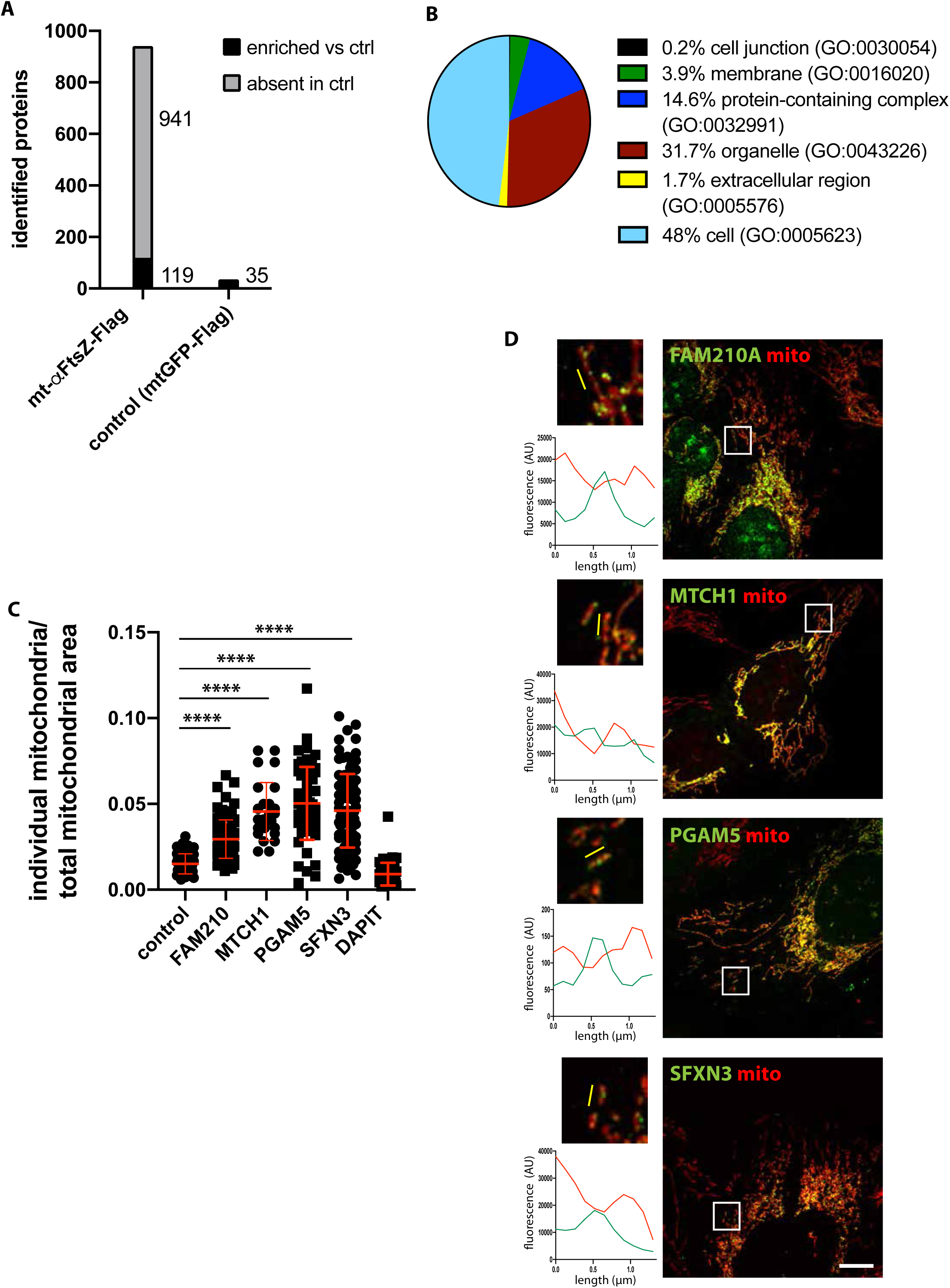
Identification of mt-αFtsZ interaction partners that play a role in mitochondrial fission. **A** Number of proteins obtained by mass-spectrometry analysis of mt-αFtsZ-Flag versus mtGFP-Flag immunoprecipitates. Grey indicates proteins identified in mt-αFtsZ-Flag and absent in mtGFP-Flag immunoprecipitates, black indicates proteins that are significantly enriched in mt-αFtsZ-Flag versus mtGFP-Flag immunoprecipitates. **B** GO-term analysis of proteins identified through mt-αFtsZ-Flag immunoprecipitatation (Panther software). **C** Semiautomated quantification (MiNA plugin) of mitochondrial morphology expressed by the amount of individual mitochondria/mitochondrial area in U2OS cells transfected with mtGFP or inner membrane proteins PGAM5, FAM210A, MTCH1, SFXN3 and DAPIT. Three experiments were pooled. Mean and SEM are displayed in red. P<0.0001 by one-way Anova, adjusted P-values < 0.0001. **D** U2OS cells expressing very low amounts of fission-inducing constructs PGAM5, FAM210A, MTCH1, SFXN3, shown in green. Mitochondria are shown in red (Hsp60). Scalebar: 10µm, insets enlarged 4x. Linescans (yellow) show varying levels of accumulation at constrictions.

We employed Flag-tagged versions to confirm mitochondrial localization of the five selected candidates and assess their impact on mitochondrial morphology. Morphometric analysis (MiNa) revealed that PGAM5, MTCH1, FAM210A and SFXN3 induced significant mitochondrial fission (Fig 4C, suppl Fig 4B). This phenotype was not a by-product of non-specific inner membrane perturbation due to overexpression, as even strong overexpression of DAPIT had no detectable effect on mitochondrial morphology. At very low expression levels, flag-tagged PGAM5, MTCH1, FAM210A and SFXN3 were also found at mitochondrial matrix constrictions (Fig 4D), supporting a possible role in matrix fission. Attempts to follow the sub-mitochondrial localization of these candidates in live cells using GFP fusions failed because their overexpression induced mitochondrial fission or substantially mislocalized to the cytoplasm (not shown). Silencing of the individual proteins did not robustly induce mitochondrial hyperfusion, and did not prevent localization of mt-aFtsZ to mitochondrial constrictions (not shown), suggesting possible functional redundancy. Co-silencing of two or more proteins resulted in high toxicity (not shown).

In conclusion, we propose that PGAM5, MTCH1, FAM210A and SFXN3 represent novel candidate effectors of inner mitochondrial membrane fission.

## Discussion

Fission of the mitochondrial matrix in the absence of outer membrane fission has been shown in metazoans, both *in vivo*^24^ and *in cellulo*^23,25,26^. How this is achieved is unclear. Although in several protozoa orthologs of the key bacterial fission protein FtsZ have been shown to localize to the mitochondrial matrix and participate to mitochondrial division^12,19,20^, FtsZ orthologs have not been detected in metazoans and fungi^10,27^. Here, we show that when directed into human mitochondria, alphaproteobacterial FtsZ (mt-αFtsZ) localizes at mitochondrial fission sites and stimulates mitochondrial division. Interestingly, expression of mitochondrial FtsZ from the brown alga *Mallomonas splendens* in yeast was found to affect mitochondrial morphology^19^. Together, this data suggest that human and yeast mitochondria contain a matrix-localized fission machinery that is structurally similar to the bacterial division machinery, as it can accommodate heterologously expressed FtsZ. indicating that mt-aFtsZ has retained features that allow it to interact productively with the inner fission machinery of today’s mitochondria.

Surprisingly, when we compared FtsZ from an alphaproteobacterium (mt-αFtsZ) with that from a gammaproteobacterium (mt-γFtsZ), only mt-αFtsZ formed stable assemblies that localize at mitochondrial fission sites. Polymerization is not unexpected *per se*, as purified FtsZ has been shown to spontaneously assemble into polymers upon GTP addition^57^. One possibility is that mt-γFtsZ polymers are unstable, as they are unable to interact with the mitochondrial matrix fission machinery, which manifests by lack of localization to mitochondrial constrictions. Another possibility is that, coming from *E.coli,* mt-γFtsZ has adapted to the diameter of the bacterium, (∼0.5µm^58^) and therefore cannot form rings in mitochondria due to spatial constrains imposed by the narrower diameter of mitochondria (∼0.2µm^59^). In contrast mt-αFtsZ may have an inherent ability to adapt to smaller diameters, as Rickettsiae have diameters as small as 0,1µm^60^.

### How are the sites of mt-αFtsZ assembly determined?

The question of how the division site is defined is central also in bacterial division. In *E. coli*, the best-studied model, two negative regulatory mechanisms have been described, based on nucleoid occlusion or on the Min system^61^. No orthologs for either of these systems have been detected in mitochondria^10^, where the nucleoid has been suggested to act as a spatial organizer of the mitochondrial fission machinery. Consistent with this view, mitochondrial fission and mtDNA dynamics are tightly linked in the red alga *Cyanidioschizon merolae*^62^. In mammalian cells, up to 70% of all fission events have been found to occur in the vicinity of a nucleoid^41^ and components of the outer (cytosolic) fission machinery have been suggested to sense the localization and replication status of nucleoids^39^. In our hands the localization of mt-αFtsZ and mitochondrial morphology were not affected by blocking mtDNA replication (Fig 3C), suggesting that this process is not essential for IFM assembly and localization. Interestingly, in nucleoid-free *E. coli* maxicells FtsZ localized to the midcell in a Min system - dependent manner^63^. As the Min system is not conserved in human mitochondria^10^, how mt-αFtsZ localizes to fission sites in the absence of nucleoids remains an open question. One possibility is that it is the inner mitochondrial fission machinery and its associated proteins, such as the constituents of contact sites or possibly lipid microdomains, that define mtDNA localization; this situation is similar to what has been observed for the mtDNA helicase Twinkle, which can associate with the inner mitochondrial membrane in the absence of mtDNA^38^.

### How does mt-αFtsZ induce mitochondrial fission at the mechanistic level?

mt-αFtsZ clearly labels mitochondrial fission sites and stimulates fission, but we currently do not know how if functions and whether it displaces components of the endogenous IFM or not. In bacteria, FtsZ has been proposed to mediate constriction either directly^57,64,65^, or indirectly, i.e. by recruiting the peptidoglycan synthesis machinery^66–68^. Mitochondria have lost the peptidoglycan, but mt-αFtsZ might act by recruiting the lipid biosynthesis machinery. In agreement with this hypothesis, we found several proteins involved in lipid synthesis among the interactors of mt-αFtsZ. However, we cannot exclude that mt-αFtsZ acts in a more direct manner, i.e. by physically pulling on the inner membrane to promote fission of the matrix compartment. The interactions that allow the recruitment of FtsZ to its membrane anchors FtsA, ZipA or SepF are mediated by the C-terminus of the protein, which is essential for promoting fission^7^. In agreement with this, the C-terminus of mt-αFtsZ was essential to localize the protein to the mitochondrial inner membrane and its deletion abolished mitochondrial fission induction. *In vitro* experiments have shown that while FtsZ can self-assemble into contractile rings in the absence of other proteins^65^, its recruitment to the membrane requires additional proteins^52^. We employed an immunoprecipitation approach to identify mitochondrial inner membrane proteins that could mediate the recruitment of mt-αFtsZ to the inner mitochondrial membrane. With this approach, we could identify proteins that have been previously implicated in mitochondrial fission and novel potential actors.

Here, we focused on five inner membrane proteins with an unclear role in mitochondrial matrix fission: PGAM5, MTCH1, SFXN3, FAM210 and DAPIT. Overexpression of DAPIT did not affect mitochondrial morphology, indicating that overexpression does not *per se* alter mitochondrial morphology, even though the mitochondrial inner membrane is one of the most protein-rich membranes^69^. In contrast, mild overexpression of PGAM5, MTCH1, SFXN3 and FAM210 induced mitochondrial fission. MTCH1 and FAM210 were not previously known to affect mitochondrial dynamics. In our hands, even mild SFXN3 overexpression induced mitochondrial fission, but previous data indicates that double deletion of SFXN3 and SFXN1 in Jurkat cells caused a decrease mitochondrial length^56^. While further studies are needed to untangle the precise role of SFXN3 in mitochondrial dynamics, our data confirms a role of PGAM5 in mitochondrial fission and adds FAM210 and MTCH1 to the growing list of inner membrane proteins playing a role in mitochondrial fission.

It is intriguing to note that among the interactors of mt-αFtsZ we also found several proteins that have been proposed to link the inner and outer membranes. One is ATAD3B, a paralog of ATAD3A, an AAA+ ATPase shown to control mitochondrial dynamics and to interact with both the inner and outer mitochondrial membranes^70^. We also found the MICOS complex component Mic60, which has been shown to bind to the nucleoid^71^ and link it with two outer membrane components (the SAM complex and Metaxins 1 and 2) and with the cytosolic fission machinery^45,46^. Strikingly, we detected SAMM50 and Metaxin 2 in mt-aFtsZ immunoprecipitates, suggesting at least partial integrity of the complex and providing a mechanism for the observed spatiotemporal coordination of mt-αFtsZ and Drp1 recruitment.

### Is mt-αFtsZ regulated?

Our live cell imaging experiments showed mt-αFtsZ localization at fission sites prior to Drp1. While we detected mt-αFtsZ at virtually all fission events, not all mt-αFtsZ assemblies underwent fission in a given time frame. This suggests that either only a subset of mt-αFtsZ oligomers are functional, or that the IFM is preassembled and poised to act upon specific triggering signals, similar to the cytosolic fission machinery based on Drp1^72^. A triggering signal for the IFM may be local calcium influx at ER-mitochondrial contact sites. Indeed, calcium influx has been shown to induce mitochondrial matrix fission, followed by outer membrane abscission^26^. Consistent with a role of calcium in the regulation of mitochondrial matrix fission, we found several regulators of mitochondrial calcium influx among the proteins that co-immunoprecipitated with mt-αFtsZ. Incidentally, divalent cations (including calcium) also stimulate FtsZ assembly and bundling in bacteria, pointing to an evolutionarily conserved regulatory mechanism of division^7,73^.

In conclusion, our work suggests the presence of a mitochondrial fission machinery in the mitochondrial matrix that retained sufficient structural conservation to accommodate a heterologously expressed bacterial FtsZ. To our knowledge, this represents the first *bona fide* marker of matrix fission sites described to date and paves the way for the molecular characterization of the mitochondrial matrix fission machinery. Future experiments will address whether FtsZ from other bacterial species have retained the ability to consistently label mitochondrial fission sites and interact with the inner mitochondrial fission machinery. Our system may thus also provide a novel “evolutionary cell biology” approach to understand which bacteria represent the closest extant relatives of mitochondria and shed light on the debated origin of the bacterial ancestor of mitochondria.

## Material and Methods

### Cloning

To create humanized versions of gammaproteobacterial FtsZ, the *E.coli* FtsZ sequence was completely re-coded according to human codon usage to optimize expression in human cells. To comply with local regulations, for alphaproteobacterial FtsZ, we used the *R.typhii* sequence as to re-code the N-terminal part (aa 2-326) of FtsZ, and the *R.prowazekii* sequence to re-code the C-terminal domain. For both constructs, we deleted the initial methionine to prevent internal initiation and the terminal stop codon was replaced with a glycine-serine linker to allow in-frame expression of a Flag or a GFP tag. Re-coded sequences were synthesized by Genecust or as gBlocks (Integrated DNA Technologies). OM-mRuby was generated by in-frame fusion of the first 215bp of human TOM20 (gBlock, Integrated DNA Technologies) with mRuby. All constructs are listed in table M1.

**Table M1:**
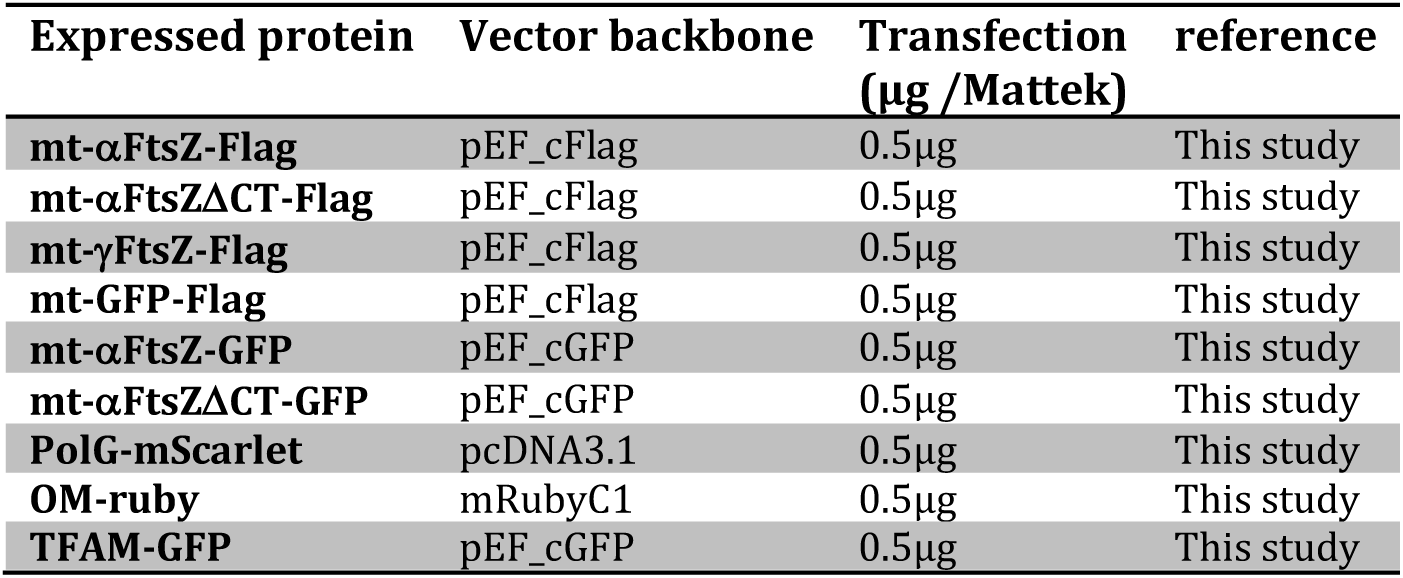
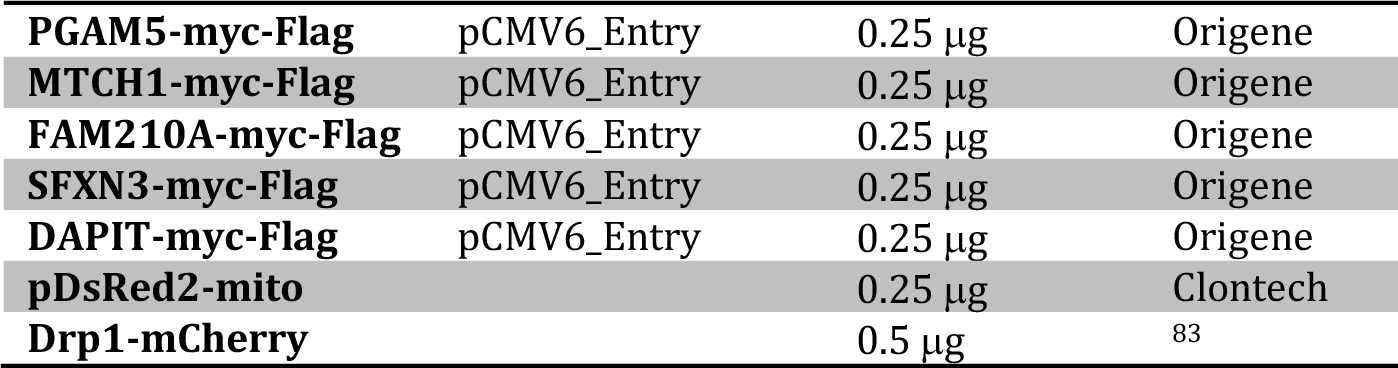
Plasmids.

### Reagents

#### Chemicals

Orange and Deep Red Mitotracker and secondary antibodies were purchased from Thermo Fisher. All chemicals were obtained from Sigma-Aldrich/Merck. Complete mini EDTA-free protease inhibitor and PhoStop phosphatase inhibitor tablets were from Roche, anti-Flag M2 dynabeads were from Sigma-Aldrich/Merck.

#### Antibodies

Antibody sources are detailed in table M2. All antibodies were used according to the manufacturer’s instructions unless otherwise stated.

**Table M2:**
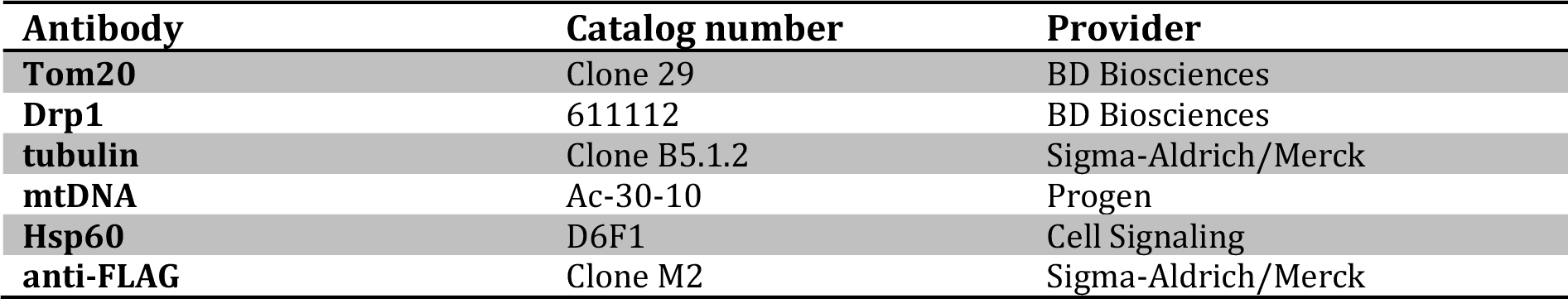
Antibodies.

| **Antibody** | **Catalog number** | **Provider** |
| --- | --- | --- |
| **Tom20** | Clone 29 | BD Biosciences |
| **Drp1** | 611112 | BD Biosciences |
| **tubulin** | Clone B5.1.2 | Sigma-Aldrich/Merck |
| **mtDNA** | Ac-30-10 | Progen |
| **Hsp60** | D6F1 | Cell Signaling |
| **anti-FLAG** | Clone M2 | Sigma-Aldrich/Merck |

### Cell culture and transfection

HeLa and U2OS cells were obtained from ATCC and cultured under standard conditions; media and additives were from Thermo Fisher. Cells were seeded on coverslips (#1.5, Marienfeld) for immunofluorescence or MatTek dishes (MatTek Corporation) for live cell imaging and transfected with FugeneHD (Roche) according to the manufacturer’s instructions, at a DNA:transfectant rate of 1:3. The DNA quantities employed for each construct are indicated in table M1. Cells were imaged or fixed and processed for immunofluorescence 24h-36h posttransfection.

### Immunofluorescence and imaging

Cells grown on coverslips were stained with mitotracker when necessary, fixed for 10 minutes in 4% paraformaldehyde (Electron Microscopy Sciences)/PBS, washed in PBS and permeabilized for 5 minutes in 0.1% Triton X-100 in PBS and blocked for at least 30 minutes in 1% BSA and 10% goat serum. Primary antibodies (see table M2) were incubated for 60 minutes in blocking buffer, followed by three washes in PBS (5 minutes) and incubation with Hoechst 33258 and Alexa-labelled secondary antibodies (Thermo Fisher) in blocking buffer for 30 minutes. Coverslips were then washed extensively in PBS and mounted in Vectashield (Vector Laboratories). For live cell imaging, cells grown on Mattek dishes were stained with mitotracker when necessary, then imaged in Fluorobrite medium (Thermo Fisher) on a Roper spinning disk confocal system (Zeiss AxioObserver.Z1 inverted fluorescence microscope equipped with an Evolve EM-CCD camera (Photometrics) and a Yokogawa CSU-X1 spinning disc). Images were acquired at 37°C with a 100x NA 1.4 oil objective using MetaMorph. Cells were imaged every 20 seconds for 10 or 15 minutes.

### Image analysis

All images were analyzed in ImageJ/Fiji (National Institutes of Health), including adjustment of brightness and contrast. Overlays were assembled in Photoshop (Adobe) and figure panels in Illustrator (Adobe). The ImageJ macro Mitochondrial Network Analysis (MiNA) toolset^74^ was used to examine mitochondrial morphology. Single cells were selected as regions of interest and pre-processing parameters were adjusted in order to obtain optimal skeletonized images of the mitochondrial network, followed by extraction of mitochondrial network features such as number of individuals, branch length and mitochondrial area, referred to as “mitochondrial area”. To obtain a degree of mitochondrial fragmentation that would be independent of the size of the mitochondrial network we used the ratio of individual mitochondria and total mitochondrial area.

### Statistical analysis

Results are expressed as means of at least three independent experiments, error bars represent the standard error of the mean. For multiple comparisons, data were analyzed with the Prism software (Graphpad) by one-way ANOVA, followed by Dunnett’s multiple comparisons test to obtain the adjusted P value. Significance is indicated as p<0.05 (*), p<0.01 (**) and p<0.005 (***), ns for p>0.05. N refers to the number of independent experiments, n refers to the number of counted events (cells, individual mitochondria, fissions).

### Electron microscopy

#### Correlative light electron microscopy (CLEM)

To obtain landmarks for CLEM the pattern of an HF-15 finder grid (AGAR) was evaporated with carbon on sapphire (3 mm diameter, 0.16 mm thickness, Wohlwendt instruments) as described^75^. The discs with stabilized carbon pattern were sterilized by UV and coated with poly-L-Lysine (Sigma-Aldrich) for cell culture. After transfection, cells were imaged in medium containing 1mM Hepes after placing the disc in a glass bottom dish (MatTek) using a Leica SP5 confocal microscope (Leica) and low expressing cells were selected. After imaging the cells were frozen with an HPM 010 (Abra fluid). Samples were freeze substituted in a Leica AFS2 (Leica microsystems) in 1% osmiumtetroxide, 0.1% uranylacetate, 5% water, 2% MeOH in dry acetone with the following schedule: 1h at - 90°C, 2.5°C/h for 16h, 30 min at −50°C, 15°C/h for 2h, 30 min at −20°C, 10°C/h for 2h, and 1h at 0°C. After substitution the dishes were infiltrated with epoxy resin (Agar) and polymerization was done in flat bottom beam capsules at 60°C for 48h. After detachment of the discs the sample was sectioned with a Leica UCT microtome (Leica microsystems) with a nominal feed of 70 nm. Sections were picked up with slot grids and contrasted with 4% aqueous uranylacetate (Merck) and Leynolds lead citrate (Delta microscopies). Images were taken with a Tecnai G2 microscope operated at 120 kV (Thermofisher), equipped with an ultrascan 4000 CCD (Gatan Inc.).

#### Immuno electron microscopy

For immune-labeling on thawed cryo sections cells were fixed with 2% PFA (EMS) + 0.1% glutaraldehyde (Sigma) in PHEM buffer, pH 7 (60 mM Pipes, 25 mM Hepes, 10 mM EGTA, 2 mM MgCl_2_) for 1 h at RT. Afterwards free aldehyde groups were quenched with 50 mM NH_4_Cl in PBS and cells were removed from the culture plastic with a rubber policeman and pelleted in a 1.5 ml eppendorf tube. The cell pellet was embedded in 12% gelatin (TAAB) and after solidification on ice, small cubes of 1 mm^3^ were cut and infiltrated overnight at 4°C with 2.3 M sucrose in PBS. The next day the cubes were mounted on metal pins and frozen by immersion into liquid nitrogen. Thin sections of 60 nm nominal feed were cut with a Leica UC6/FC6 cryo-microtome at −120°C. The sections were picked up with a 1:1 mixture of 2% methylcellulose in water and 2.3M sucrose in PBS. After thawing the sections were deposited on grids and labelled with rabbit anti GFP (Rockland) followed by Protein A gold (CMC Utrecht). At the end of the labelling the sections were contrasted with 0.4% uranylacetate in 1.8% methylcellulose and airdried before observation with Tecnai G2 microscope. For the quantification of matrix/inner membrane localization of mt-αFtsZ-GFP or its ΔCT version, 36 and 25 random sections were quantified respectively.

### Immunoprecipitation

Immunoprecipitation was performed as described^76^ with modifications. Briefly, HeLa cells were seeded on 10 cm dishes and transfected with 7µg DNA. After 36h, cells were washed three times in PBS and lysed for 30 min with 1 ml lysis buffer/10cm dish (20 mM Tris, pH 7.4, 100 mM NaCl, 10% glycerol, 1,5mM MgOAc) supplemented with 0.5% NP-40 (Igepal), 1x protease and phosphatase inhibitors (Roche). Lysis and all subsequent steps were performed at 4 °C. After lysate clarification at 13000x*g* for 10 minutes, the protein concentration of the supernatant was determined by Bradford assay (Pierce). 1 mg of lysate was incubated overnight with 20µl anti-Flag M2 magnetic Dynabeads (Sigma-Aldrich/Merck) under shaking. Magnetic beads were recovered, washed three times with lysis buffer and four times with washing buffer (50 mM Tris, pH 7.4, 150 mM NaCl) and eluted with 2×20µl 3xFlag peptide (100 mg/mL in washing buffer). The experiment was performed in triplicate. For western blot, 5µl (10%) eluate was supplemented with 2x Laemmli buffer, boiled for 10min resolved on a gradient SDS-PAGE (Biorad), and subjected to western blotting via wet transfer to 0.45µm nitrocellulose membrane (Millipore). Ten µg total lysate were loaded (corresponding to 1%) for the input.

### Proteomic analysis

#### Protein digestion

Proteins were solubilized in urea 8 M, NH_4_HCO_3_ 50 mM pH 7.5, then disulfide bonds were reduced with 5 mM tris (2-carboxyethyl) phosphine (TCEP) for 30 min at 23°C and alkylated with 20 mM iodoacetamide for 30 min at room temperature in the dark. Samples were diluted to 1 M urea with 50 mM NH_4_HCO_3_ pH 7.5, and Sequencing Grade Modified Trypsin (Promega, Madison, WI, USA) was added to the sample at a ratio of 50:1(w/w) of protein to enzyme for 8 h at 37°C. Proteolysis was stopped by adding 1% formic acid. Resulting peptides were desalted using Sep-Pak SPE cartridge (Waters) according to manufactures instructions. Peptides elution was done using a 50% acetonitrile (ACN), 0.1% FA buffer. Eluted peptides were lyophilized and then store until use.

#### LC-MS/MS analysis

a nanochromatographic system (Proxeon EASY-nLC 1000, Thermo Fisher Scientific) was coupled online to a Q Exactive–Plus Mass Spectrometer (Thermo Fisher Scientific). For each sample, 1µg of peptides was injected onto a 50 - cm homemade C18 column (1.9µm particles, 100 Å pore size, ReproSil-Pur Basic C18, Dr. Maisch GmbH) and separated with a multi-step gradient from 2% to 45% ACN at a flow rate of 250 nl/min over 180 min. The column temperature was set to 60°C. The data were acquired as previously described^77^.

#### Data processing

Raw data were analyzed using MaxQuant software version 1.5.3.8^78^ with database search parameters as described in^77^. The MS/MS spectra were searched against Uniprot proteome database of *Human (January 13, 2015, 20,432 entries)* and mt-αFtsZ-Flag protein, and usual MS contaminants. Data were quantified with the MaxLFQ algorithm by requiring a minimum peptide ratio count of 2. The parameter “match between run” was checked. Raw data have been deposited to the ProteomeXchange Consortium via the PRIDE^79^ repository with the dataset identifier PXD016722.

#### Statistical and functional analysis

For the statistical analysis of one condition versus another, proteins exhibiting fewer than 2 intensities in at least one condition were first discarded. After log2 transformation, intensities were normalized by median centering within conditions (*normalizeD* function of the R package DAPAR^80^). Proteins without any intensity in one condition (quantitatively present in a condition, absent in another) were considered as differentially abundant. Next, missing values were imputed using the *imp.norm* function of the R package *norm*. Proteins with a log2(fold-change) inferior to 1 have been considered as proteins with no significant difference in abundance. Statistical testing of the remaining proteins was conducted using a limma t-test^81^. An adaptive Benjamini-Hochberg procedure was applied on the resulting p-values to select a set of significantly differentially abundant proteins with a false discovery rate of 1%^82^. The proteins of interest are therefore the proteins that emerge from this statistical test supplemented by those being quantitatively absent from one condition and present in another. Gene ontology analysis of mass spectrometry results was performed with the online softwares Panther (http://pantherdb.org/) and DAVID (https://david.ncifcrf.gov/), using standard settings. Transmembrane proteins were predicted using the online software TMHMM (http://www.cbs.dtu.dk/services/TMHMM/).

## Data availability

The data that support the findings of this study are available from the corresponding author on reasonable request.

## Author contributions

AS: Data curation, Formal analysis, Investigation, Methodology; MS: Investigation, Methodology, Writing – review & editing; TNT: Investigation, Methodology; MM: Methodology; FS and PC: Funding acquisition, Writing – review & editing; FS: Conceptualization, Data curation, Formal analysis, Investigation, Methodology, Validation, Visualization, Writing – original draft, Supervision.

## Acknowledgements

We would like to thank Véronique Hourdel and Quentin Giai Gianetto for initial analysis of mass spectrometry data and Ludmila Bonnand for help with live cell imaging analysis. Alessandro Pagliuso, Jan Riemer and Tim Wai are thanked for discussion and Simonetta Gribaldo, Bastian Huelsmann, Nika Pende, and Hans Spelbrink for critical reading of the manuscript. This study was supported by the European Research Council (H2020-ERC-2014-ADG 670823-BacCellEpi to P.C.) and Institut Pasteur. A.S. was supported by a BioSPC doctoral fellowship from the Université Paris Diderot. P.C. is a Senior International Research Scholar of the Howard Hughes Medical Institute. F.S. is a CNRS permanent researcher.

## Supplementary Figure legends

**Suppl Fig1:**
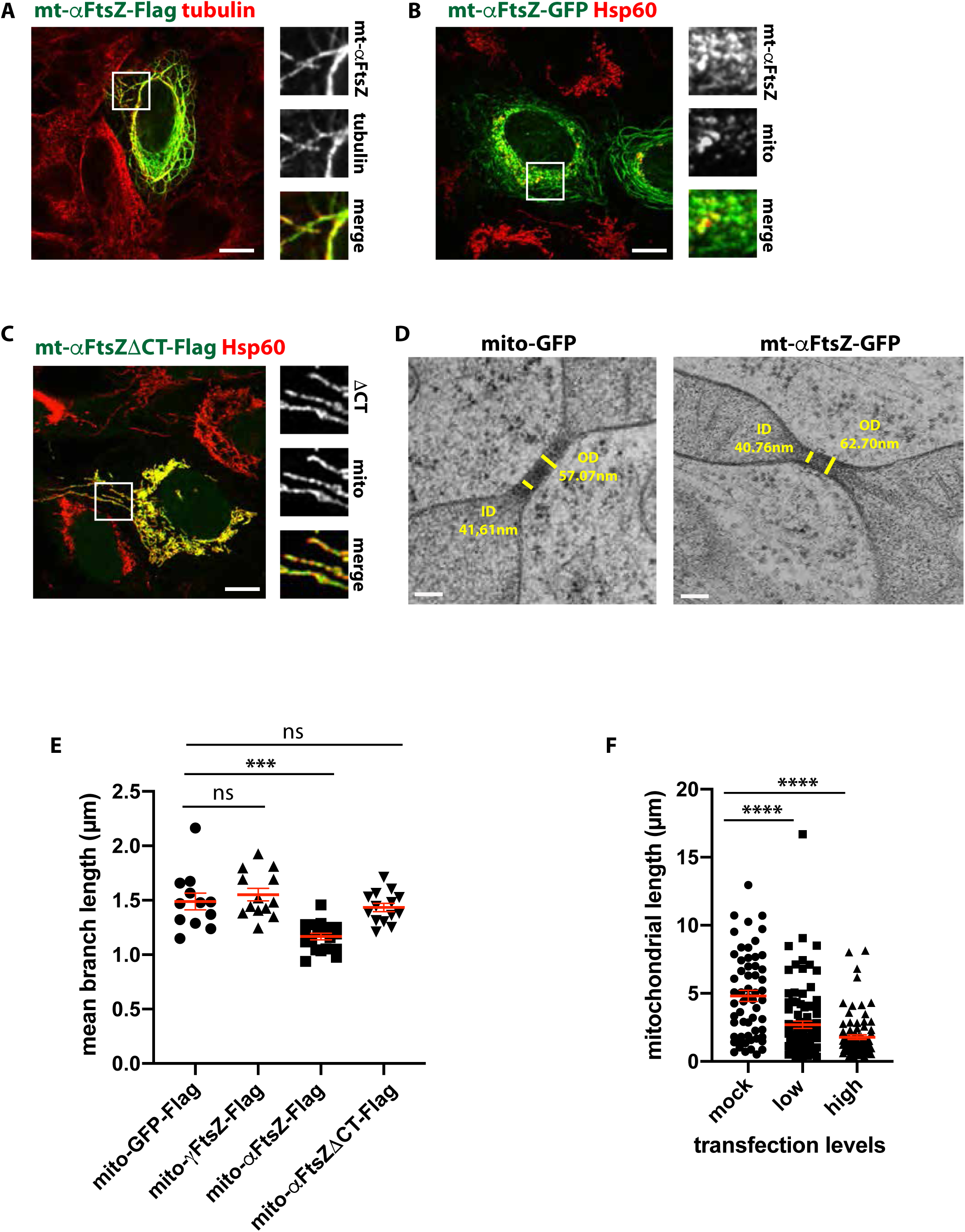
Characterization of mt-αFtsZ localization and their impact on mitochondrial morphology. **A** Cytosolic mt-αFtsZ-Flag (green) colocalizes with tubulin (red). Scalebar: 10µm, insets enlarged 3x. **B** Perinuclear aggregation of mitochondria in cells where mt-αFtsZ-GFP mislocalizes to the cytosol. Immunofluorescence of U2OS cells transiently expressing mt-αFtsZ-GFP (green) and labeled for Hsp60 (mitochondria, red). The percentage of cells displaying cytosolic filaments increased to 20.2±10.5% (n=1361, N=4) when GFP was employed instead of the flag tag. Scalebar: 10µm, insets enlarged 3x. **C** Representative example of a cell displaying diffuse mt-αFtsZΔCT-Flag (green) staining in mitochondria. Mitochondria are shown in red (Hsp60). Scalebar: 10µm, insets enlarged 3x. **D** Mitochondrial constrictions in Hela cells expressing intermediate levels of mt-αFtsZ-GFP or mt-GFP analyzed by HPF-CLEM. Inner diameters (ID) and outer diameters (OD) are indicated in yellow. Scalebar: 100nm. **E** MiNA analysis of mitochondrial branch length in U2OS cells transfected with mito-GFP-Flag or mitochondrially targeted FtsZ from a gamma- (mt-γFtsZ-Flag) or alphaproteobacterium (mt-αFtsZ-Flag, C-terminal deletion mutant mt-αFtsZΔCT-Flag). The same dataset was used as in Fig 1C. Mean and SEM are displayed in red, p<0.0001 by one-way Anova, adjusted P-value <0.0001. **F** Manual measurement of the length of resolvable mitochondria in HeLa cells expressing mt-αFtsZ-Flag and counterstained for mitochondria (Hsp60). n>60, N=5. Mean and SEM are shown in red, P<0.0001 by one-way Anova, adjusted P-values <0.0001.

**Suppl Fig 2:**
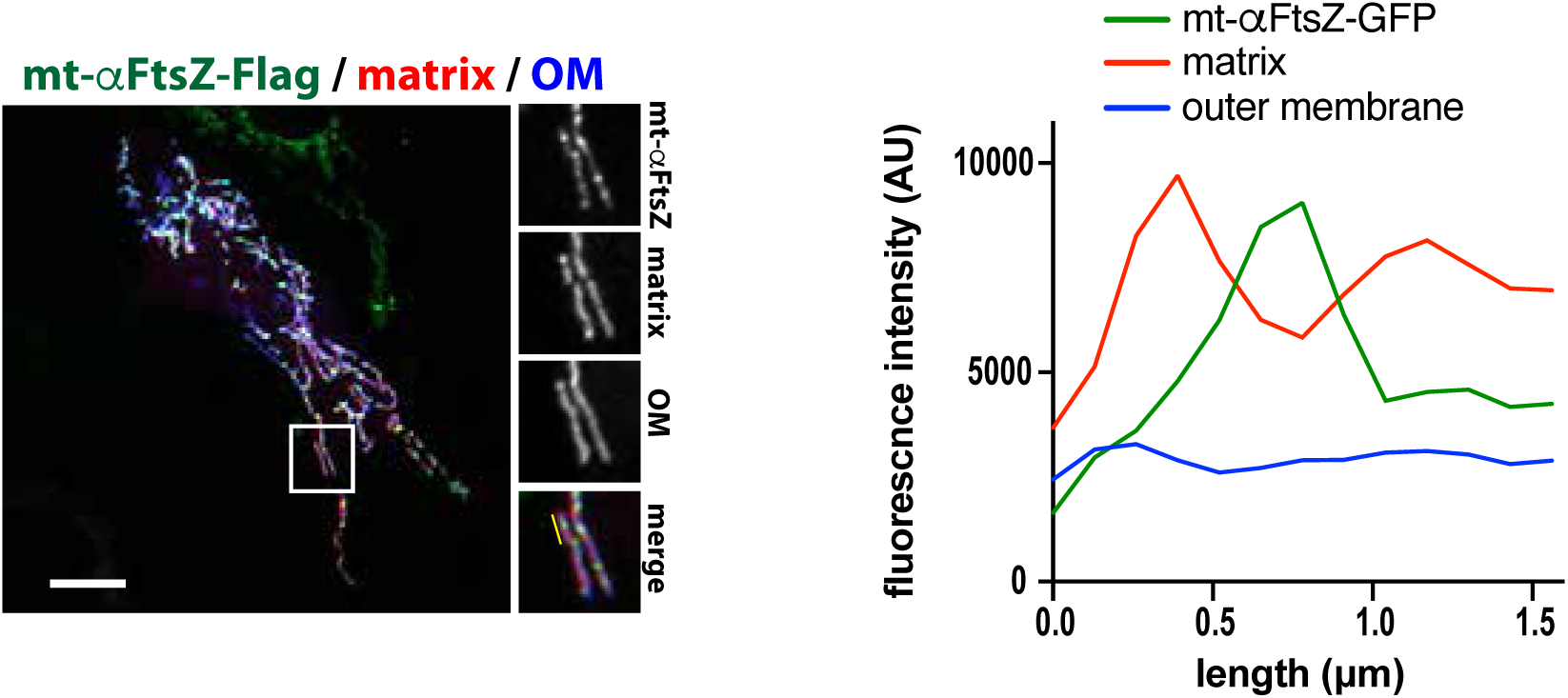
Differential effect of mt-αFtsZ on the outer membrane and the mitochondrial matrix. HeLa cells labeled for mt-αFtsZ-Flag (green) display matrix constriction (Hsp60, red) in the absence of outer membrane (Tom20, blue) constriction

**Suppl Fig 3:**
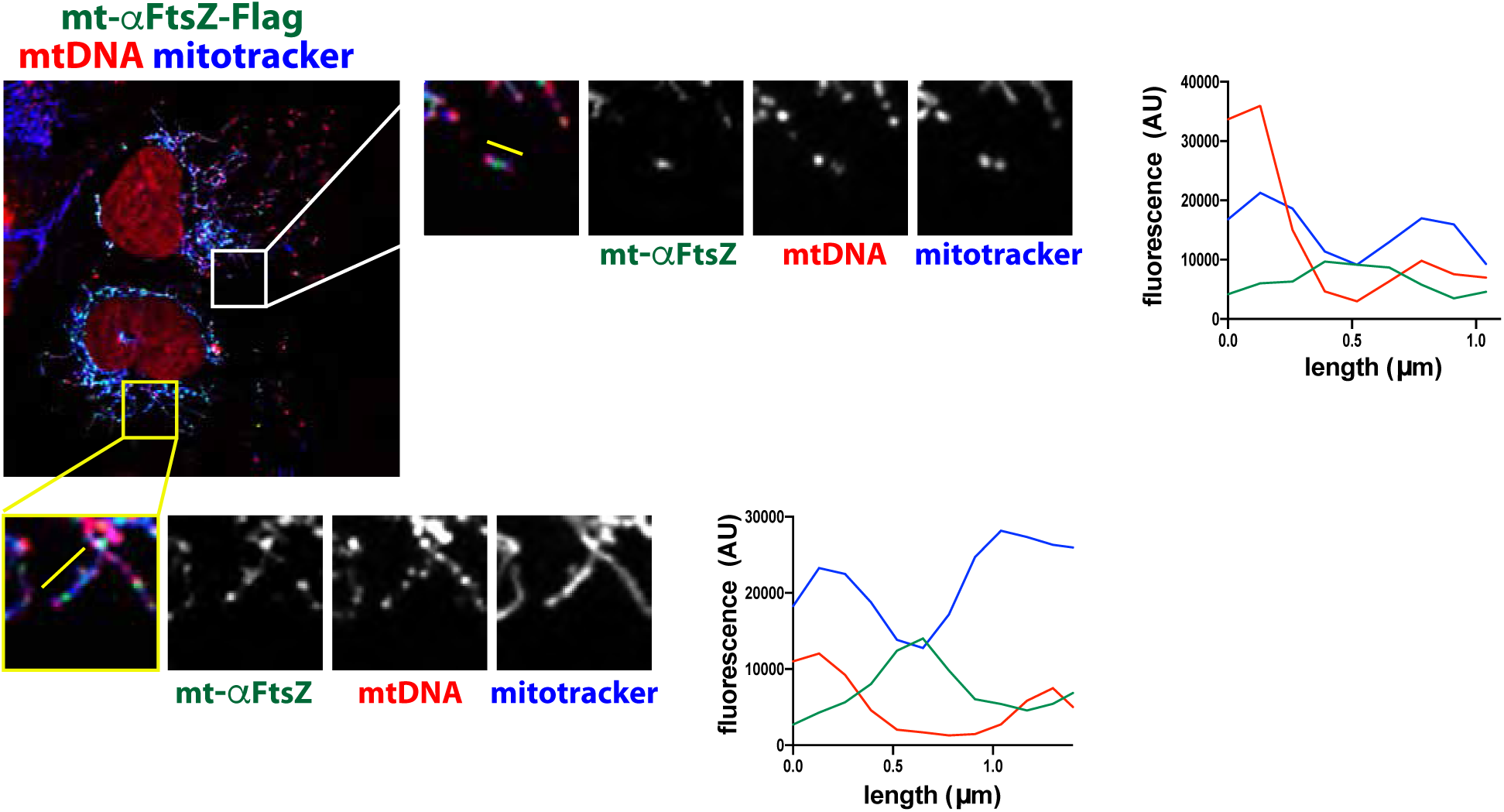
mt-αFtsZ-labeled constrictions are flanked by mtDNA. U2OS cells transfected with mt-αFtsZ-GFP (green) and labeled for mtDNA (red) and mitotracker deep red (blue). Insets enlarged 2x. Linescans showing mt-αFtsZ-GFP localization at mitochondrial constrictions, flanked by mtDNA.

**Suppl Fig 4:**
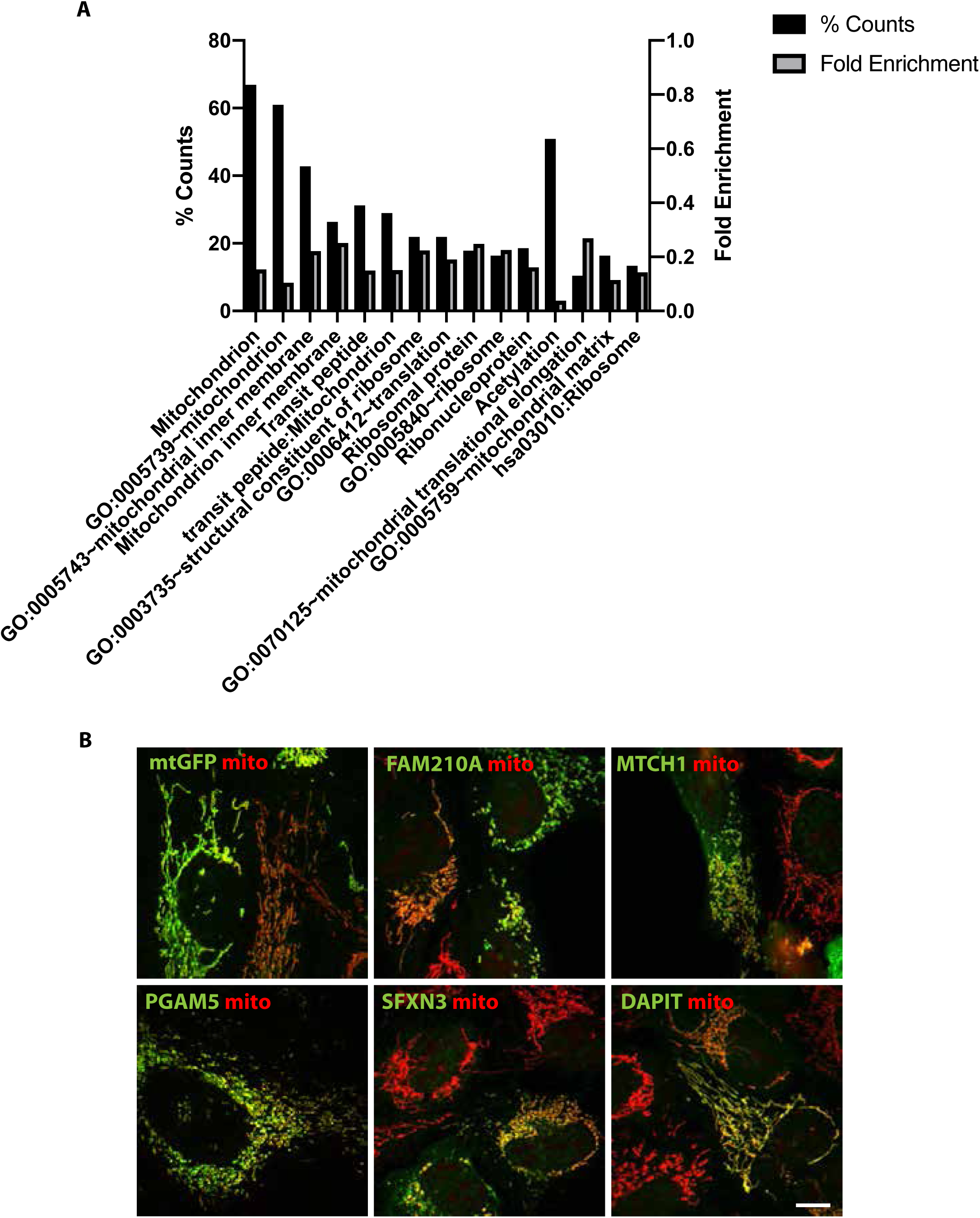
Functional annotation of mt-αFtsZ interaction partners and overexpression of selected proteins. **A** Functional annotation clusters (“Biological Process”, BP) of proteins co-immunoprecipitating with mt-αFtsZ-Flag assessed in the background of the human proteome (DAVID software). The 4 most enriched clusters are shown, with associated keywords and GO terms. Black bars show % protein count and grey bars show fold enrichment. Benjamini scores are shown on the far right. All p-values were<0.01. **B** Representative images of U2OS cells transfected with flag-tagged versions of the inner membrane proteins FAM210A, MTCH1, PGAM5, SFXN3 and DAPIT (shown in green) and quantified in Fig 4D. Mitochondria were labeled with Hsp60 (red) and appear fragmented by FAM210A, MTCH1, PGAM5 or SFXN3 expression.

